# Ribosome profiling reveals extensive translational alterations in steatohepatitis-associated hepatocellular carcinoma

**DOI:** 10.64898/2026.02.11.705272

**Authors:** Asier González, Meric Ataman, Aleksei Mironov, Niels Schlusser, Muskan Pandey, Alexander Schmidt, Mairene Coto-Llerena, Salvatore Piscuoglio, Nitish Mittal, Mihaela Zavolan

## Abstract

Hepatocellular carcinoma (HCC) is one the most lethal cancer types. While infections with hepatitis B or hepatitis C viruses remain the primary risk factors for HCC development worldwide, metabolic disease-associated steatohepatitis has become the fastest growing etiology of HCC, particularly in the West. Our study characterizes, for the first time, the remodeling of the translation landscape in steatohepatitis-associated HCC using ribosome profiling and mass spectrometry techniques. In striking contrast with the transcriptional alterations, which do not show strong dependence on associated co-morbidity, translational alterations are broader in the steatohepatitis compared to the viral background. These alterations affect hundreds of genes, whose products are often involved in subcellular protein localization. By quantifying the relationship between the ribosome occupancy of upstream or downstream open reading frames (uORFs or dORFs) and coding regions (CDSs), we provide evidence of uORFs attenuating the translation efficiency of CDSs in HCC. We also identify numerous novel translated regions, including from RNAs currently annotated as non-coding, some yielding peptides that can be reproducibly identified in mass spectrometry datasets. Our study thus provides novel molecular data on HCC with a focus on the steatohepatitis etiology, reveals unexpectedly extensive translational control in this co-morbidity, and gives multiple hints and candidate targets for follow-up experiments.

## Introduction

Liver cancer is the fifth most common cancer world-wide and the third leading cause of cancer-related deaths. It arises in the background of chronic liver disease, caused by infection with hepatitis B or C viruses, and increasingly more often by metabolic dysfunction in the context of alcohol use, diabetes or obesity [1,2]. The main type of liver cancer is hepatocellular carcinoma (HCC). Genomic [3,4], proteomic [5], and integrative multi-omic [6–8] analyses have been carried out to decipher the molecular alterations occurring in this heterogeneous disease and enable more specific treatments. However, the progression from individual types of chronic liver diseases to HCC remains poorly understood.

Liver has one of the highest rates of protein synthesis among adult mammalian organs [9–11], regulated during all its essential biological activities: insulin-dependent fatty acid synthesis, lipogenesis, glycolysis, ATP production [12,13]. Interest in how mRNA translation is remodeled to support the survival and proliferation of cancer cells is growing [14–17], as translational control could enable cancer cells to rapidly adapt to stress without relying on new mRNA transcription. Various mechanisms of translational control could play a role in HCC – such as high expression of eIF4E or high PI3K-AKT-mTOR signaling [18–20] – but their investigation in human tissue samples has remained limited.

A technique broadly used to quantitatively study the translation dynamics with subcodon resolution is ribosome profiling/footprinting [21], also known as ribo-seq. It has generated a wealth of data primarily from cell lines [22], with data from human tissues starting to emerge [23]. In addition to canonical protein-coding sequences (CDSs), recent studies have uncovered a variety of non-canonical open reading frames (ncORFs), some generating peptides essential for cancer cell survival and proliferation [18,24,25], others giving rise to neoantigens [26,27]. NcORFs can be located upstream or downstream of the main CDS (uORFs and dORFs, respectively), or in transcripts currently annotated as non-coding (long non-coding RNA, lncRNA). Miscoding and stop codon readthrough arising from perturbed expression of small nucleolar RNAs [19] or from nonstop extension mutations [28,29], can further diversify the proteome of cancer cells. Although ribosome profiling has been applied to hepatitis B virus (HBV)-associated HCC to reveal limited translational control [30], no such analysis has so far been done for metabolic dysfunction-associated HCC.

In this study, we carried out ribosome profiling and mass spectrometry-based measurements of protein expression in 43 samples from 24 patients to uncover the translation landscape of HCC with various associated co-morbidities. We further integrated transcriptome data from The Cancer Genome Atlas (TCGA, [4]), to assess the relative contribution of transcriptional and translational control in HCC occurring on the background of viral infection or non-viral liver disease. Our data reveals more extensive translational control of gene expression in HCC with non-viral co-morbidity, mediated in part by uORFs. We further found evidence of translation for numerous long non-coding RNAs (lncRNAs). Our study provides an extensive resource for studies of translational control in HCC.

## Results

### Ribosome profiling of human liver samples

We obtained data from an etiologically heterogeneous cohort of 24 patients clinically classified before the recent change in nomenclature [31]: 11 had steatohepatitis (SH) of unknown – alcoholic or non-alcoholic – origin (ASH/NASH), 3 had ASH, 3 were hepatitis B virus (HBV)-positive, 2 were hepatitis C virus (HCV)-positive, 1 had ASH/NASH and was also HCV-positive, and the remaining 4 had unclear co-morbidity. In 17 of the 24 cases we analyzed tumor (T) and matched non-tumoral tissue (NT) from the same patients, and in one case, three distinct tumor sites were sampled (Fig. 1A, Suppl. Table 1). Following the experimental design shown in Fig. 1A, we obtained ribosome profiling libraries of 18-28 million reads per sample, with read lengths peaking at 30-31 nucleotides (Fig. 1B and Suppl. Table 2). Upon mapping the reads to the transcriptome (see Methods), we found that 30-65% of reads in a library were derived from mRNAs (Suppl. Table 2), with 85-95% of these mapping to annotated CDSs (Fig. 1C). We inferred the location of the P site in each read using our previously established approach [32,33] (see Methods) and observed that 40-60% of reads in a library matched the annotated codon frame (Fig. 1D). Altogether, these metrics fulfill the expected quality criteria for ribosome profiling data [34], notably given that resected tissue samples often yield RNA of lower concentration and integrity compared to cell lines [23].

**Figure 1.**
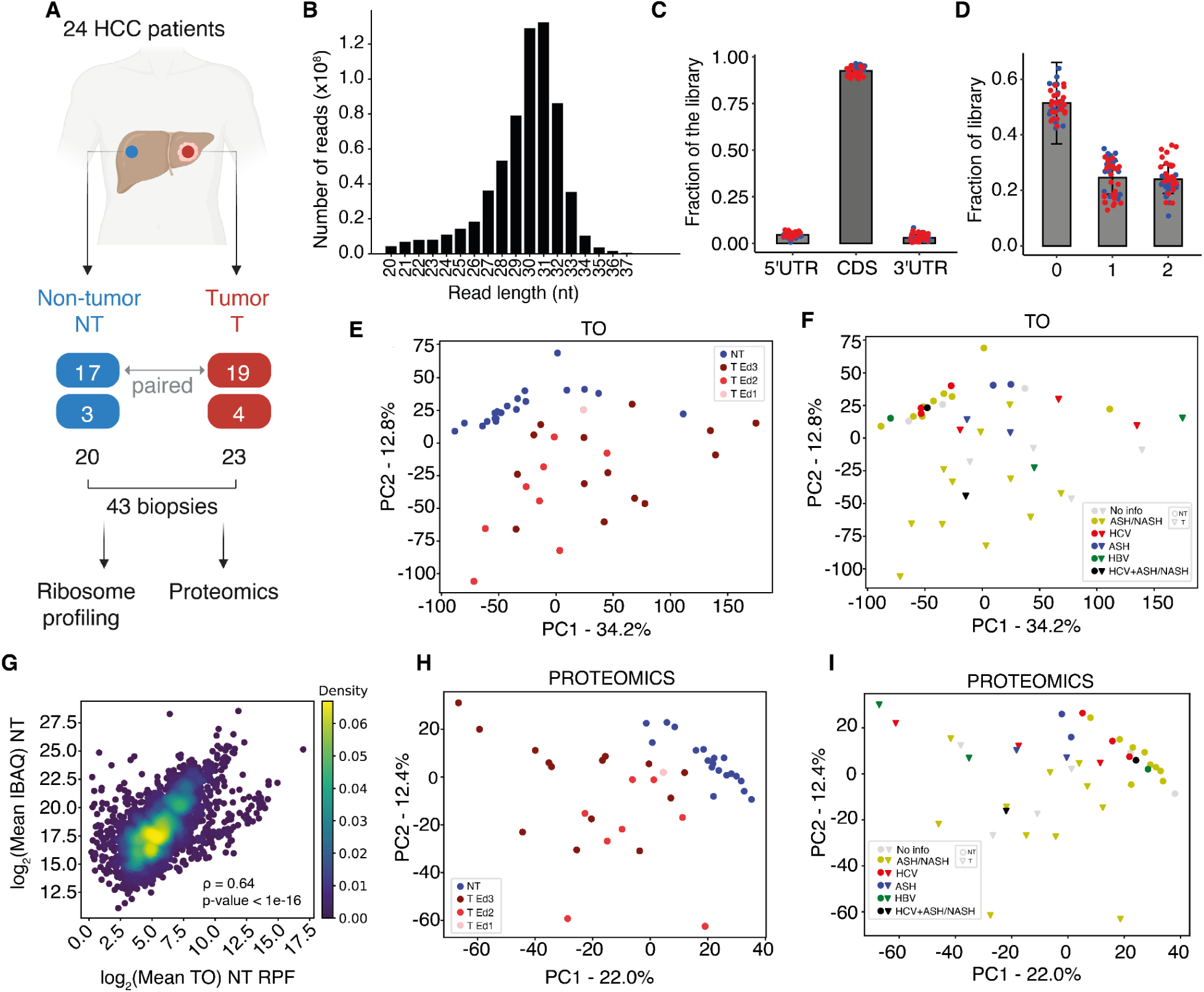
Summary of the data set. **A.** Experimental design: 43 biopsies from 24 HCC patients were analyzed by ribosome profiling, including 16 tumor and matched non-tumoral (T-NT) pairs, one case of 3 tumors - 1 non-tumor samples, 3 unpaired tumors and 4 unpaired non-tumor samples. In panels A-D, samples are indicated by the red (T) and blue (NT) colors, respectively. Image created with BioRender. **B.** Read length distribution of pooled ribosome profiling libraries. **C.** Fraction of RPFs in each library mapping to 5’UTR, CDS, or 3’ UTR. **D.** Fraction of RPFs with inferred P-site mapping to each of the 3 reading frames of annotated transcripts. **E.** PCA of gene TO values calculated from ribosome profiling data. Tumor samples are shaded according to the Edmondson grade; higher Edmondson grade indicates a less differentiated tumor. **F.** Similar to E, but coloring the samples by co-morbidity and T/NT status. **G.** Correlation of sample-average TO with the sample-average protein iBAQ signal across protein-coding genes quantified in both assays, for non-tumor samples. **H.** Similar to E, based on normalized protein-level intensity values from proteomics. **I.** Similar to F, based on protein-level intensity values from proteomics.

We detected ribosome-protected fragments (RPFs, i.e. reads mapped uniquely from the procedure described above) from ∼19’800 genes, with ∼11’000 genes being represented with at least one RPF in each sample. We estimated the translational output (TO) of individual mRNA species as the normalized density of RPFs per CDS nucleotide; technically, this is the well-known metric of transcripts per million (TPM), computed from RPFs rather than RNA-seq reads. We transformed the values to log_2_ scale, standardized them, and carried out principal component (PC) analysis (PCA). The first two PCs explained almost half (47%) of the variance in TO, with tumor samples separating well from matched non-tumoral tissue (Fig. 1E, p-value = 0.001 in ClusterSignificance analysis [35]). We did not observe a further clear clustering of tumor samples by Edmonson grade (p-value = 0.111, ClusterSignificance analysis, Fig. 1E), or presence/absence of cirrhosis (p-value = 0.178, ClusterSignificance analysis, Suppl. Fig. 1A). However, we did observe significant clustering when we distinguished the tumor samples by co-morbidity backgrounds, namely viral infection and steatohepatitis (p-value = 0.0035, ClusterSignificance analysis, Fig. 1F).

We further measured the proteomes of all samples by mass spectrometry in a data-independent acquisition (DIA) manner, detecting 5’000-6’000 proteins per sample. Virtually all genes whose expression could be quantified by mass spectrometry were also quantified by ribosome profiling and the Spearman correlation coefficient of the sample-average TO with the level of the corresponding protein was 0.64 and 0.65 for non-tumor and tumor samples, respectively (Fig. 1G for NT samples, Suppl. Fig. 1B for T samples). Following standard quantification, normalization and standardization (see Methods), the first two PCs of the proteome data explained 34.4% of the variance (Fig. 1H), and the ClusterSignificance results reflected largely those from the ribosome footprints. That is, tumor and non-tumor samples separated from each other (p-value = 0.0001), with no clear separation by Edmondson grade (p-value = 0.536), or presence/absence of cirrhosis (p-value = 0.074, Suppl. Fig. 1C). In contrast to the ribosome profiling data, the viral/non-viral co-morbidity did not yield significant clustering (p-value = 0.0549), which could be due to the smaller number of genes being measured in proteomics compared to ribosome profiling.

The transcriptome-wide TO profiles were more similar when comparing non-tumor samples with each other and less similar when comparing tumor with non-tumor samples or tumor samples with each other (Suppl. Fig. 1D). The true tumor/non-tumor pairing of samples from individual patients did not yield significantly different results compared to random pairing. The proteomics measurements yielded similar results (Fig. 1I, Suppl. Fig. 1E). Of note, samples with lower Edmondson grades were on average, more similar to non-tumor ones than those with higher grades, both at the level of TOs and protein levels (Suppl. Fig. 1F-G), consistent with prior reports [8]. Altogether, these results demonstrate the consistency of ribosome profiling and proteomics measurements and highlight that malignancy has a strong effect on the translational profile of the liver, while the grade, presence or absence of cirrhosis, co-morbidity and other patient-specific features (e.g. germline genetic variation) appear to have smaller impact. The latter allowed us to conduct further analysis not accounting for matching between T-NT pairs.

We further checked the consistency of the data from our etiologically heterogeneous cohort and those previously obtained from 10 HCC patients, 9 of whom were HBV-positive [30]. We reanalyzed the latter with our pipeline and observed that for genes quantified in both datasets (17’773 of the 17’796 in the Zou study [30]) the sample-averaged TO values correlated very well when conditions were matched (NT-NT or T-T, Pearson correlation coefficients of 0.95 and 0.93, respectively, Suppl. Fig. 1H), and slightly less when tumor samples from one study were compared with non-tumor samples from the other (NT-T or T-NT, Pearson correlation coefficients of 0.89 and 0.92, Suppl. Fig. 1I). Thus, our RPF data is highly consistent with prior measurements, while almost tripling the number of available HCC ribosome profiles by providing so-far lacking information on the non-viral co-morbidity. Consequently, the functional categories of genes that are up- or down-regulated at TO level in HCC relative to non-tumor tissue are also similar in the two datasets (Suppl. Fig. 1J). We further observed the previously reported suppression of the urea cycle and arginine-to-polyamine conversion enzymes, alongside the upregulation of downstream polyamine metabolism genes in HCC [36] (Suppl. Fig. 1K). Finally, the T vs. NT protein level changes that we measured matched well those reported in the study of Ng et al. [8] (Suppl. Fig. 1L). Thus, our ribosome profiling data recovers broad patterns of gene expression changes expected for HCC, providing us with the basis for investigating more specific questions about the translational control of this disease, in particular with respect to the viral and non-viral co-morbidities.

### Quantification of transcriptional and translational control in HCC

The number of ribosome footprints captured for an individual gene (and the derived TO) is determined by both the abundance of the mRNA and by the efficiency with which ribosomes are recruited and translate the mRNA. The latter quantity is typically called translation efficiency (TE), and is estimated as the ratio of the library-size-normalized RPF counts to the library-size-normalized counts of RNA-seq reads, both mapped to mRNA coding regions [21]. To distinguish which of these factors underlie the observed TO changes, measurements of mRNA abundance are needed. While the limited amount of material we obtained in our study did not allow us to carry out both ribosome profiling and RNA-seq from the same samples, a large HCC RNA-seq dataset is available from TCGA [4], with associated metadata, including co-morbidity. To assess the suitability of this dataset for our analyses, we checked its consistency with the data from the above-mentioned study of Zou et al. [30], where ribosome profiling and RNA-seq data were obtained from the same 10 tissue samples. T-NT changes in RNA levels in virus-infection-associated HCC samples from TCGA were moderately correlated (Pearson r = 0.63, Suppl. Fig. 2A) with those measured by Zou et al. [30], possibly due to a reduced RNA integrity in Zou et al. samples relative to TCGA, as indicated by transcript integrity scores [37] (Suppl. Fig. 2B). Nevertheless, the T-NT changes in TO estimated from the Zou et al. ribosome profiling data correlated equally well with the changes of mRNA levels estimated either from the same samples or from the TCGA dataset (Pearson r = 0.72 vs. 0.74, Suppl. Fig. 2C-D). This indicates that TCGA provides a suitable – and, at the same time more comprehensive – reference of mRNA level expression in HCC, which we could use along with our TO data to identify translation-level effects.

To first characterize the transcriptome remodeling in HCC we submitted the TCGA RNA-seq data to the Integrated Motif Activity Response Analysis (ISMARA) [38]. We found an increased activity of cell proliferation-related transcription factors (TFs) in T samples relative to NT controls, while the activity of TFs linked to metabolic and detoxification functions is reduced (Fig. 2A). In particular the YBX1, FOS, NFYC, NFYA, NFYB, and CEBPZ TFs acting on binding sites with the core motif ATTGG formed a central hub of factors with increased activity in tumors, stimulated by MYC, and promoting the activity of E2F family members (Fig. 2B). These results are in line with reports of MYC and E2F integrating several HCC-relevant upstream oncogenic pathways [39–41] to implement a switch from specialized liver functions to cell proliferation. Importantly, no significant difference in TF activities was detected between tumors of viral or non-viral origin.

**Figure 2.**
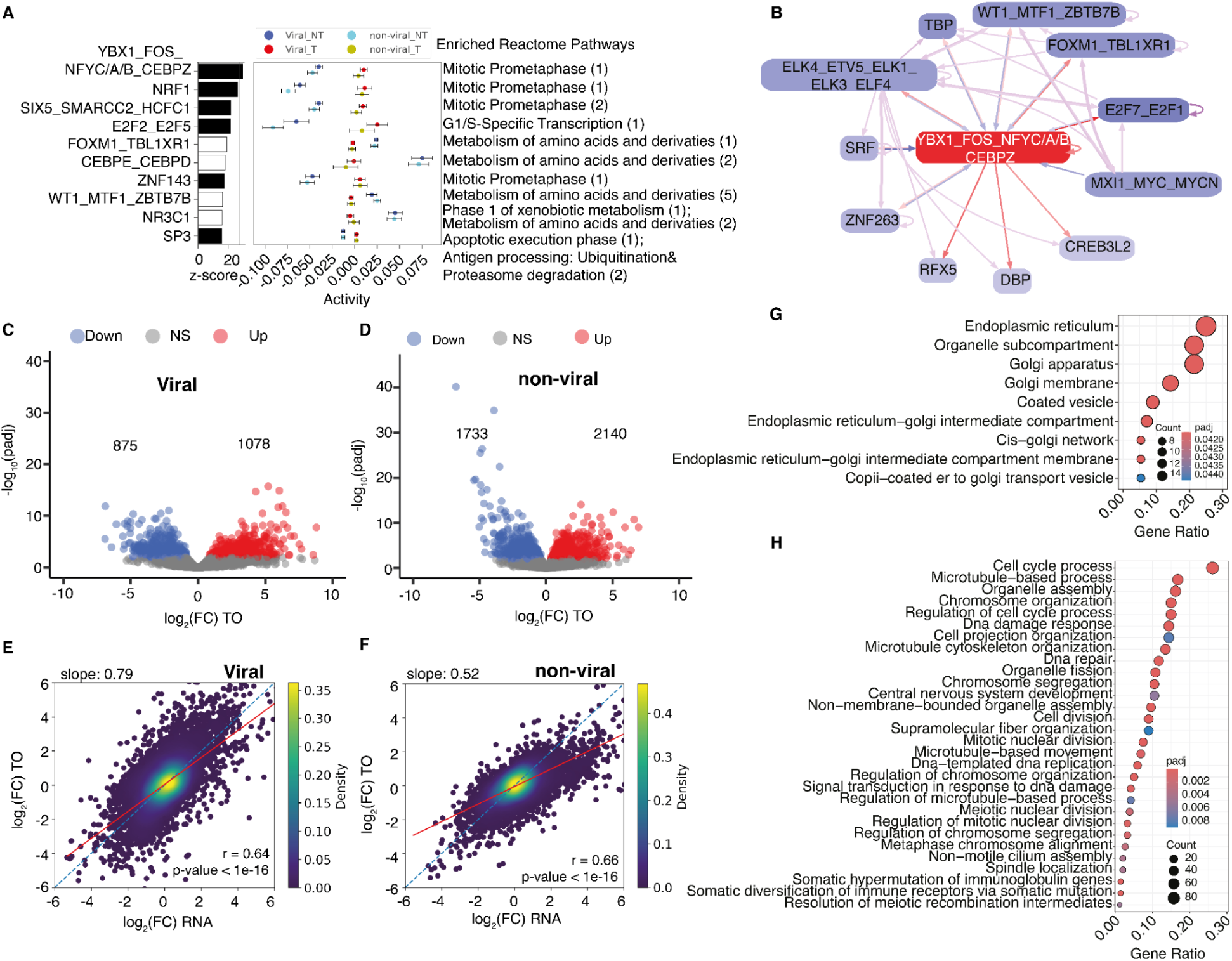
Transcriptional vs. translational control of gene expression in HCC. **A.** ISMARA results for top 10 transcription factor binding site motifs with strongest explanatory power in TCGA RNA-seq data, as assessed by their Z-values (left-most part of the panel) for both disease backgrounds. The center part of the panel depicts average motif activities in the tumor (red) and non-tumor (blue) samples. Reactome [44] enrichment analysis of each motif’s target genes was obtained from ISMARA detailed report, obtained categories were further ranked by significance (total log-likelihood). Selected categories and their ranks are shown in the right part of the panel. **B.** First level transcriptional regulatory network centered on the YBX1-FOS-NFYC-NFYA-NFYB-CEBPZ TFs. Outgoing regulatory interactions of the central factors are shown with red arrows, incoming ones with blue, and in violet are interactions not involving the central factors. **C.** Changes in the translational output of mRNAs in tumor (T) relative to non-tumor (NT) samples for viral background (cutoff at adjusted p-value < 0.05 from DESeq2 differential analysis). **D.** Same as in panel C, for non-viral samples. **E.** Relationship of mRNA level changes in T vs. NT samples from the TCGA RNA-seq dataset (x-axis), and RPF level changes in our ribosome profiling dataset for viral samples. The diagonal line (light blue, dashed) corresponds to changes in translational output that are entirely explained by changes in mRNA levels. Paired ordinary least squares (OLS) regression line is shown in red, and its slope is depicted at the top-left of the panel. **F.** Same as in panel E, for non-viral samples. **G.** Cellular Component (CC) GO terms overrepresentation analysis for genes that have upregulated TE in non-viral samples. For the GO term analysis in panels G and H, the cutoff for the adjusted p-value was 0.05. **H.** Biological processes (BP) GO terms overrepresentation analysis for genes that have downregulated TE in non-viral samples. The x-axis for panel G and H represents the proportion relative to all altered genes and top 30 terms (by significance) are shown.

We then carried out a differential TO analysis separately for samples associated with either viral infection or steatohepatitis, and found a more substantial remodeling of the translatome in the latter (Fig. 2C-F), reflected in the significantly different slopes (p-value = 6.29e-83, Welch’s t-test) of pairwise linear regression models of TO change (ribosome profiling) as a function of mRNA change (TCGA RNA-seq, Fig. 2E-F). In other words, the TO changes more closely reflect mRNA changes in tumors occurring on the background of viral infection than on the background of steatohepatitis. Importantly, we obtained similarly high slopes for our virus-associated samples (slope = 0.79, Fig. 2E) and the virus-associated samples from the Zou et al. study, and in the latter case, both when the paired RNA-seq (slope = 0.77, Suppl. Fig. 2C, p-value = 0.074, Welch’s t-test for slope comparison with the viral cohort from our study) or the TCGA RNA-seq (slope = 0.84, Suppl. Fig. 2D) were used as reference for ribosome profiling. These results indicate that the difference we observed between samples of viral and non-viral background is neither due to our dataset containing relatively few virus-associated samples, nor to systematic batch effects of cross-dataset comparisons.

We further conducted differential translation efficiency analysis to identify mRNAs undergoing specific translational control [21]. Applying a previously developed method based on the DESeq2 package [42] with a custom linear model termed deltaTE [43], we identified 65 genes with increased TE and 350 with decreased TE in steatohepatitis background, both at the adjusted p-value cutoff of 0.05 (Suppl. Table 3). In the virus-associated HCC there were only 13 genes with altered TE at the same significance cutoff. Genes with increased TE tend to encode proteins involved in protein transport and localizing in the endoplasmic reticulum and Golgi (Fig. 2G). In contrast, genes related to the cell cycle and DNA replication tend to have reduced TE (Fig. 2H). Thus, our data reveals a higher prevalence of translational control in HCC of non-viral association compared to virus-associated HCC, translational changes generally “buffering” the mRNA level changes.

### Identification of uORFs potentially impacting the translation of canonical ORF

A well studied mechanism for modulating the translational output of CDSs involves uORFs (reviewed in [45–47]). While generally hindering the translation of the downstream, canonical ORFs, specific combinations of uORFs are also instrumental for the translation of mRNAs encoding proteins that help alleviate stress [48–50]. To determine whether uORFs may drive some of the changes in TE that we observed in HCC with non-viral background we first extracted translated uORFs from our samples, shown by the pattern of 3-nucleotide periodicity of RPF coverage (p-value of 0.01 in Wilcoxon’s rank sum test, Methods, Fig. 3A). With a previously developed computational workflow [32] and requiring reproducible coverage across samples (see Methods), we obtained 4’105 uORFs, the majority in the length range of ∼20-80 codons (∼2^6^-2^8^ nucleotides in Fig. 3B). Changes in the TOs of uORFs and CDSs were highly correlated across samples, reflecting the fact that they are driven by changes in the levels of mRNAs that contain both the uORFs and the corresponding CDSs. Indeed, randomizing the linkage between uORF and CDS abolished this correlation (Suppl. Fig. 3A).

**Figure 3.**
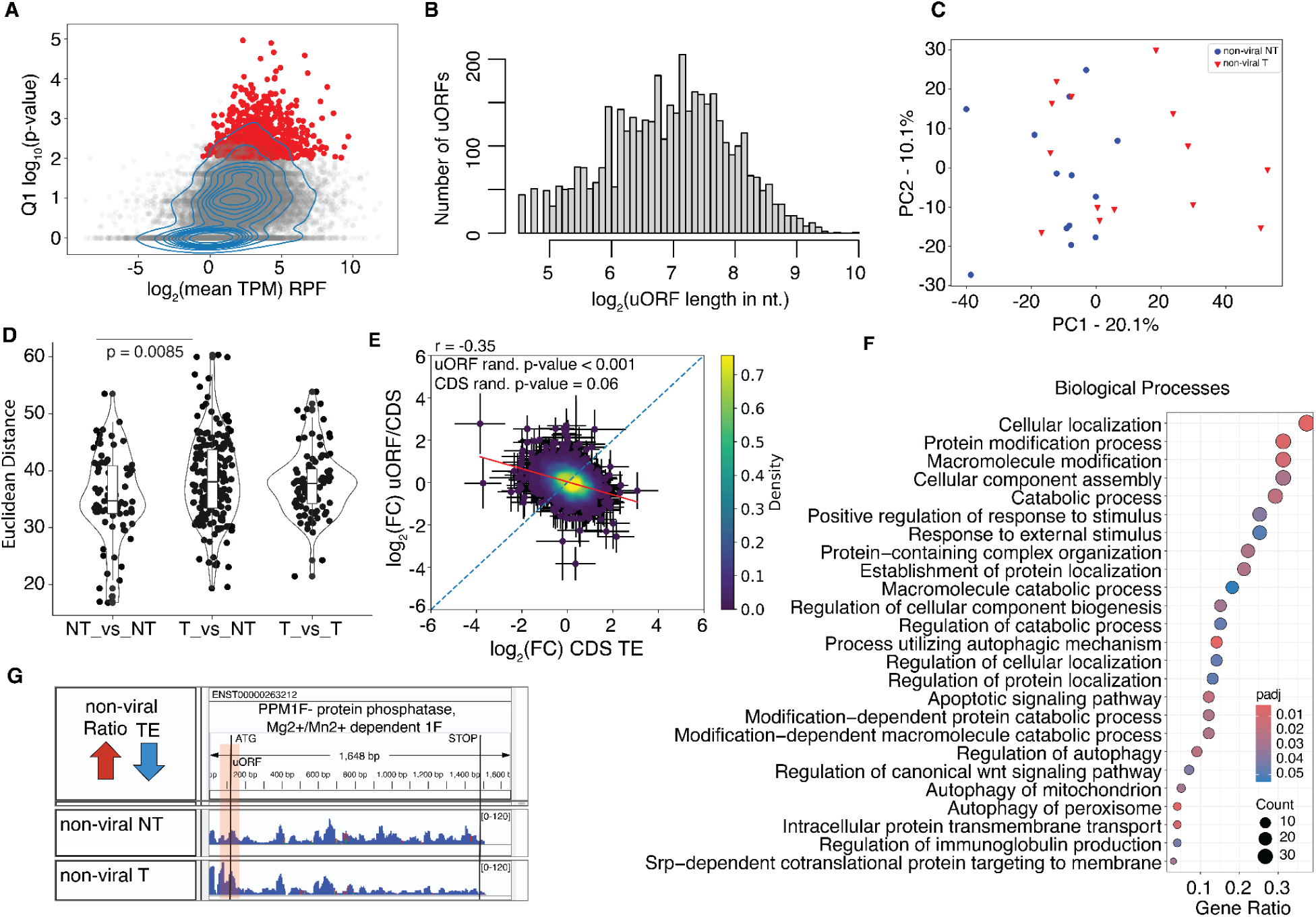
uORF-associated translational control in HCC. **A.** Selection of uORFs with evidence of translation, based on the strength of evidence of 3-nucleotide periodicity of RPFs (see Methods). Q1 represents the first quartile of p-values for all samples. **B.** uORF length distribution. Only the uORFs with significant frame bias (in red in panel A) are shown. **C.** Principal component analysis (PCA) of uORF/CDS ratios (p-value = 0.29, for separation of viral tumor samples from the non-viral tumor samples, by ClusterSignificance analysis). **D.** Euclidean distances between uORF/CDS ratios of individual genes in pairs of NT and T-NT samples from viral/non-viral disease background groups. **E.** Average log_2_ fold-change in uORF/CDS ratio across T-NT for non-viral samples shown as a function of the change in the CDS TE in the same samples. The log_2_FC in ratio was estimated by DESeq2 using a custom linear model deltaTE approach, with condition (c) component in the model as T/NT, sequencing type (s) component as uORF/CDS counts. Error bars indicate standard error estimates for the log_2_FCs according to DESeq2 analysis. **F.** Biological process GO term overrepresentation analysis of genes with significantly changed uORF/CDS ratios in samples of non-viral background. **G.** Example of an mRNA (from PPM1F gene) with anti-correlated uORF/CDS and TE changes in T vs. NT. The coverage shown in the IGV browser represents the sum of RPFs in all samples of the specified type.

To then identify candidates of uORF-mediated translational control, we assessed the evidence of RPF redistribution between uORFs and CDSs in T vs. NT samples, calculating the uORF/CDS RPF ratio for all genes harboring translated uORFs. While T and NT samples did not clearly separate on the first two PCs of the uORF/CDS ratios (Fig. 3C, p-value = 0.06, ClusterSignificance analysis), the distances between the vectors of uORF/CDS values were larger when comparing T and NT samples than when NT samples were compared with each other (Fig. 3D), indicating a degree of RPF redistribution between uORF and CDS that in tumors goes beyond the variability of non-tumor samples. At the same time, the distribution of T-NT changes in uORF/CDS RPF ratio was narrow and largely symmetrical around log_2_ values of 0 (Suppl. Fig. 3B, Suppl. Table 4), indicating that the redistribution is very limited in magnitude and the number of mRNAs potentially affected. Nevertheless, we carried out a series of tests to determine the relationship between uORF and CDS translation in the HCC system.

If uORFs had no bearing on CDS translation, the uORF/CDS RPF ratio and CDS TE should fluctuate independently across conditions. This is not what we observed. Rather, changes in uORF/CDS RPF ratio between T and NT samples were negatively correlated with the changes in CDS TE (Fig. 3E). To again exclude the linkage between uORF and CDS expression explaining the anti-correlation, we shuffled the uORF RPF counts across transcripts and repeated the uORF/CDS - CDS TE correlation analysis. In none of the randomized datasets did we obtain as low a correlation coefficient as in the real dataset (Fig. 3E, Suppl. Fig. 3C). Thus, the fact that uORFs and CDSs are co-expressed on the same mRNA does explain the uORF/CDS - CDS TE correlation. However, as we cannot measure these quantities independently, another potential confounder is the CDS RPF counts appearing in the denominator of the uORF/CDS ratio and in the numerator of the TE. To exclude this explaining the observed anti-correlation, we next randomized the CDS RPF counts between genes. The p-value of the uORF/CDS - CDS TE correlation coefficient obtained for the real dataset with respect to the randomized ones was 0.06 (Fig. 3E, Suppl. Fig. 3D). These results indicate some of the uORFs we identified do have a repressive effect on CDS translation in HCC, a mechanism that may be exploited to selectively fine-tune translation in tumors occurring on the background of metabolic disease.

However, as mentioned above, the magnitude of RPF redistribution of between uORFs and CDSs is small, with only 9 genes reaching an *adjusted* p-value cutoff of 0.05 of the change in uORF/CDS ratio (Suppl. Table 4). Upon relaxing the significance metric from adjusted to raw p-value of 0.01 and accounting for the dependency of p-values on the sample size in differential analysis by repeated sub-sampling of TCGA RNA-seq data (see Methods) we observed that out of 1’111-3’205 (depending on the resampled dataset) genes with significantly changing TE, only 0.7%-2.0% (21 genes, on average) had a significant (same raw p-value < 0.01), opposite direction change in the uORF/CDS ratio. At the same time, of the 109 genes with significant change in uORF/CDS ratio 10%-29% (depending on the resampled RNA-seq data set) had a significant, opposite direction change in TE. Thus, our data confirms the well-known repressive effect of uORFs on CDS translation and reveals tens of candidates for which uORF-dependent modulation operates in HCC occurring on the background of metabolic disease. However, most of the TE changes do not seem to involve uORF-dependent regulation.

The genes with significant increased or decreased uORF/CDS ratio changes in T vs. NT non-viral samples come from biological processes related to protein traffic, localization and, notably, autophagy (Fig. 3F). The RPF coverage along the protein phosphatase, Mg2+/Mn2+ dependent 1F (PPM1F), is shown as an example in Fig. 3G.

### Translation beyond the CDS in HCC

As ribosomes lacking specific rRNA modifications have been implicated in miscoding and stop codon readthrough in liver cancer cell lines [19], we also assessed the evidence of translation downstream of annotated stop codons in our data. With the same methodology of ORF annotation that we employed for uORFs, we found substantially fewer ORFs with evidence of translation (189) located in 3’UTRs (downstream ORFs, dORFs) compared to 5’UTRs (4’105). The lengths of translated dORFs and uORFs were similar, in the range of ∼20-80 codons (Fig. 4A). Also similarly to uORFs, changes in dORF/CDS RPF ratio were anti-correlated with the changes in CDS TE (Fig. 4B, Suppl. Fig. 4A), but in this case, the anti-correlation could be explained entirely by confounders (Suppl. Fig. 4B-C). Thus, we did not find evidence of a regulatory impact of dORFs on the CDS, though in some cases the dORF coverage is quite high, comparable to the CDS (Fig. 4C).

**Figure 4.**
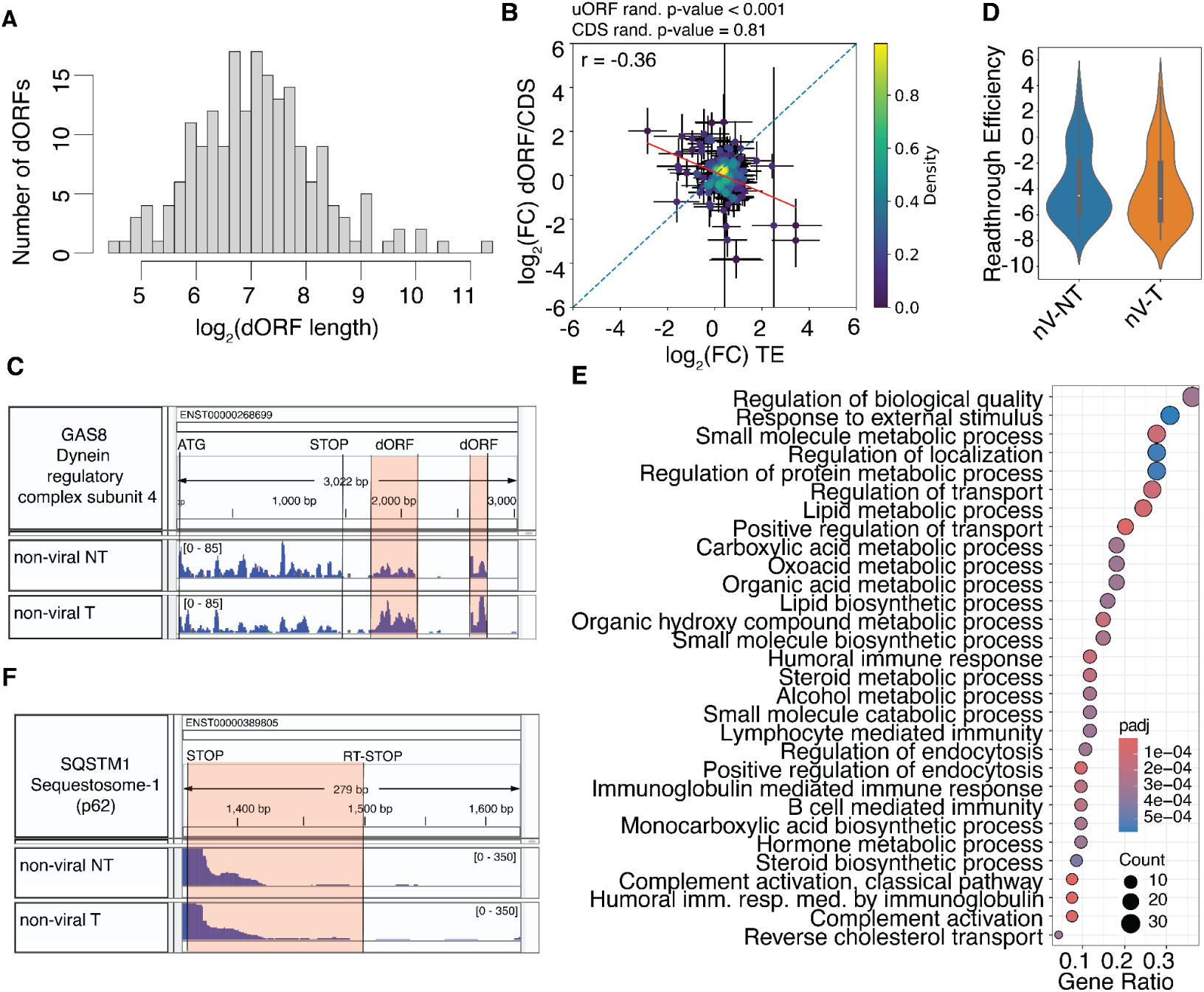
dORF/RT analysis. **A.** Length distribution of dORFs with significant frame bias. **B.** Average log_2_ fold-change in dORF/CDS ratio in T-NT samples with non-viral background shown as a function of the change in CDS TE. Error bars indicate standard error estimates for the log_2_ fold-changes according to DESeq2 analysis. **C.** Example of the GAS8 mRNA, containing in its 3’UTR dORFs (highlighted in red) with as high coverage by RPFs as the CDS. The coverage is shown in the IGV browser, and one track represents the sum of RPFs in all samples of the specified condition. **D.** RT/CDS RPF density log_2_FC distribution in T vs NT for samples of non-viral disease background. **E.** Biological processes GO term overrepresentation analysis of all genes with evidence of SCRT translation. **F.** Example of an mRNA with significantly reduced SCRT in tumor T relative to NT samples shown in the IGV browser. Indicated RT-STOP indicates the location of the SCRT-induced downstream stop codon that is significantly supported by 3-nucleotide periodicity of RPF coverage.

Prior studies also reported the generation of proteins with specific subcellular localization via stop codon readthrough (SCRT) [51,52]. Searching for evidence of SCRT in our data, we identified 113 genes with significant frame preference of RPFs mapping downstream of the stop codon and robust coverage in all samples of a given condition (T/NT from non-viral background). The RPF density in the SCRT region relative to the CDS is approximately -5 on the log_2_ scale, suggesting that ∼3% of proteins synthesized from these mRNAs carry C-terminal extensions, similar in T and NT conditions (Fig. 4D). SCRT proteins come from functional categories related to response to stimuli and immunity, and they include endocytosis, complement activation, lipid metabolism, other metabolic functions and endomembrane system (Fig. 4E, Suppl. Fig. 4D). The RPF coverage around the stop codon of the previously identified potential readthrough target Sequestosome-1 (SQSTM1) [53] is shown in Fig. 4F. SQSTM1 SCRT isoform has also mass-spectrometry evidence from multiple independent proteomics datasets (Suppl. Fig. 5, Suppl. Table 5). We observed a decrease in SCRT efficiency in tumors (Suppl. Figure 4E, Suppl. Table 4), and to exclude that the bias for decreased SCRT efficiency is due to systematic changes in the proportions of immune cells in T relative to NT samples, we estimated the proportions of various cell types from expression data with standard methods. We did not find significant or consistent results (Suppl. Fig. 4F-M), suggesting that the cell type composition is not sufficient to explain the observed differences. However, it is possible that the observed reduction in SCRT reflects an increased senescence of immune cells in tumors, as reported before [32,54,55].

### Upregulation of lncRNA translation in HCC of non-viral background

Long noncoding RNAs (lncRNAs) have been strongly linked to HCC, largely through transcriptional effects (reviewed in [56]). However, an increasing number of studies have identified lncRNAs that undergo translation and generate functional peptides [57,58]. To check the presence of such lncRNAs in our system, we mapped the reads that did not match rRNAs or mRNAs to the lncRNA set from the version 6 of NONCODE database [59] and then evaluated the evidence of 3-nucleotide periodicity of ribosome footprints on lncRNA-encoded ORFs. With the same criteria as for other types of translation units (significant reading frame preference and robust expression with at least 1 RPF in all samples of a condition, viral/non-viral NT or T), we identified 1’842 translated lncRNA-derived ORFs. Their length distribution was shifted slightly towards larger values relative to lncRNA-encoded ORFs that did not pass our criteria of translation evidence (Fig. 5A, B). The RNA level changes of these lncRNAs in the virus and non-virus-associated samples were highly correlated (Fig. 5C), yet we observed the same trend of reduced slope of TO vs. RNA level changes in the non-viral HCC (Fig. 5D,E), as for CDSs. The translation efficiency of lncRNA-derived ORFs was decreased in virus-associated HCC and increased in the non-viral background (Fig. 5F), with some lncRNAs already reported to promote HCC progression – ST8SIA6 antisense RNA 1, SNHG11, CYTOR (Fig. 5G) and HCP5 [60,61]^,[62],[62,63]^ – being significantly upregulated. An HCP5-derived micropeptide has also been implicated in ferroptosis in gastric cancer [64]. 168 of the 1’842 candidates are supported by at least one spectrum in the proteomics data from our study or from the 16 liver-related datasets from the PRIDE database [65] (Suppl. Table 5). The lncRNAs with most peptide evidence from multiple samples are shown in Fig. 5H. They include lncRNAs with increased TO in tumor samples from non-viral disease background, some already linked to cancer. In particular, the translated ncORF of ZFAS1 was reported to promote cancer cell migration by increasing reactive oxygen species in HCC [66] and SPATA41 has been related to breast cancer [67], though no peptide evidence has been reported for it so far. Thus, our data provides further support for lncRNA-derived peptides, suggesting a more prominent contribution of these peptides in non-viral associated HCC.

**Figure 5.**
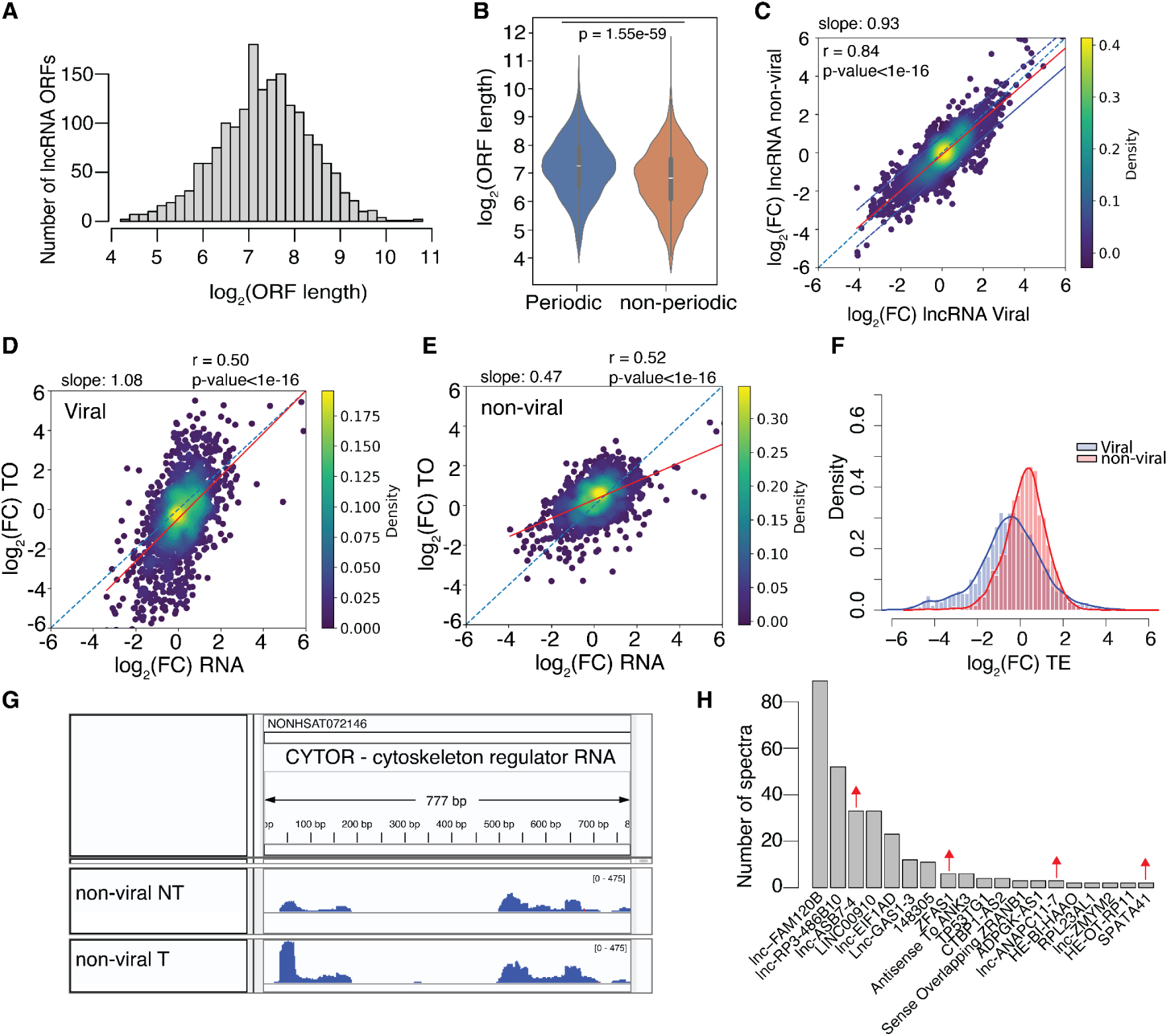
Analysis of lncRNA translation. **A.** Length distribution of lncRNA-derived ORF with significant frame bias. **B.** Box plots showing the length distribution of predicted ORFs in lncRNA with (blue) and without (red) evidence of 3 nt-periodicity in our ribosome profiling data. **C.** 2D density plot of average log_2_FC (T vs NT) in RNA levels of lncRNAs in samples from viral (x-axis) vs. non-viral background (y-axis). **D.** Correlation of RNA level changes (T vs. NT samples of viral background) in samples from the TCGA dataset (x-axis) with RPF level changes in our dataset for lncRNA-derived ORFs. The diagonal line (orange, dashed) corresponds to changes in translational output that are entirely explained by changes in mRNA levels. **E.** Same as panel F for samples of non-viral background. **F.** Distribution of TE changes (T vs. NT) of lncRNA-derived ORFs in samples from viral and non-viral background. **G.** RPF coverage of lncRNA CYTOR, shown in the IGV browser. **H.** Proteomics evidence from multiple datasets for lncRNA-derived ORFs. Red arrows indicate lncRNAs that are upregulated at RPF level in T vs. NT for the non-viral background.

Altogether, our study presents a first assessment of translational control in HCC with non-viral, steatohepatitis background, revealing a more prominent role of this regulatory layer in metabolic disease relative to viral infection-associated HCC. In particular, we identified uORFs whose RPF coverage is anti-correlated with the TE of the corresponding CDS and 12 lncRNAs whose translation efficiency is significantly upregulated (adjusted p-value<0.05, according to deltaTE method) in metabolic disease-associated HCC.

## Discussion

HCC accounts for approximately 80% of liver cancer cases [68] and is the third leading cause of cancer-related death worldwide [1,69]. Prior studies have shown that the proteogenomic profile of HCC is quite heterogeneous despite mutations focused on the same targets: the coding regions of TP53 and CTNNB1 genes, and the promoter of TERT [8]. It is known that oncogenic processes related to translational control are perturbed in HCC [8,30], but the implications for gene expression remain poorly understood.

The rate-limiting step of translation is initiation [70], regulated by the context-dependent activation of translation initiation factors (eIFs) interacting with regulatory elements in 5’UTRs, particularly uORFs [71]. Several eIFs have been linked to liver pathophysiology [10,72–74], including the eIF4F complex [75,76], eIF5A [77,78], eIF6 [12,13], and eIF2A [79]. To unravel the extent, targets and mechanisms of translational control in HCC, we have carried out deep sequencing of ribosome footprints from a substantial cohort of matched T-NT samples. We focused primarily on HCC with non-viral background, for which data have so far been unavailable [30]. Our samples came from 24 patients, most of whom were diagnosed with metabolic dysfunction-associated steatohepatitis, while 5 were positive for HBV or HCV.

By further integrating mRNA-seq data from TCGA [80], we were able to distinguish between transcription and translation-level effects. Transcriptional alterations were largely similar in viral-and steatohepatitis-associated HCC, entailing a reduction in liver-specific metabolic functions and upregulation of proliferation-related activities mediated by the same set of transcription factors (Fig. 2). Strikingly, while in the viral-associated HCC the translational output of mRNAs was mostly determined by mRNA levels, as reported before [30], this was not the case for steatohepatitis-associated HCC. In the latter, hundreds of genes had significantly altered translation efficiency suggesting a prominent role of translational control, which we investigated further. Interestingly, changes in transcript levels were buffered at the level of translation, as was recently reported in multiple organisms [81,82].

Upstream ORFs are an important lever of translational control, trapping ribosomes to repress the translation of downstream CDSs. Many studies have demonstrated the relevance of uORF-mediated translational control during stress, when mRNAs encoding stress response proteins escape this repression and undergo translation, despite a general decrease in protein production [83]. uORFs hinder the 5’UTR scanning by the small ribosome subunit, and may also result in the activation of the nonsense-mediated decay pathway [83–85]. Data from massively parallel reporter assays also demonstrated the general, repressive effect of uORFs on mRNA translation [86,87], while large-scale genotyping studies found that uORF-creating variants are under negative selection, similar to missense variants in the CDS region [88,89]. Ribosome-profiling-based studies have uncovered a wealth of uORFs [90], but only few of these have been functionally characterized. In particular, whether uORFs modulate CDS translation in a nuanced manner, outside of strong perturbations that lead to a stress response, is debated [83,91].

As the tumor microenvironment provides various stresses, potentially rendering the uORF-dependent translational control more relevant, we sought to characterize the relationship between uORF and CDS translation in HCC. In viral infection-associated HCC we found only limited evidence of translational control, in line with a prior study that studied a larger cohort of virus-associated HCC [30]. However, the non-viral background showed a distinctly different picture, genes associated with membrane-delimited compartments and protein subcellular localization exhibiting increased TE (Fig. 2G). Redistribution of ribosome footprints between uORFs and CDSs may be involved in a small proportion (1-2%) of the significant TE changes (Fig. 3E, Suppl. Fig. 3), in line with results obtained in yeast [91]. This percentage is likely underestimated, as suggested by the frequent upregulation of uORF-containing mRNAs upon depletion of RNA decay factors [92,93]. As the repression of CDS translation can lead to RNA decay [94], the chance of capture of ribosome footprints from repressed transcripts to infer their uORF/CDS redistribution may be reduced. However, approximately a quarter of the mRNAs with significant redistribution of RPFs between uORFs and CDSs undergoes a significant change in CDS TE, indicating that uORFs act as attenuators of CDS translation specifically in disease contexts such as HCC. At the same time, for the majority of translated uORFs we could not find evidence for a translation-regulatory role, and their functions remain to be determined. Some of these uORFs yield peptides detectable in multiple datasets from both liver tissue and cell lines (Suppl. Fig. 5), and they may carry out independent functions.

The coupling between uORF and CDS translation is actively investigated [95,96]. A recent study reported that the reinitiation of translation downstream of ATF4 uORFs upon stress requires the DENR-MCTS1 complex or the homologous EIF2D protein [97], and EIF2D. Interestingly, while many translation initiation factors were upregulated in our HCC data relative to non-tumor samples, DENR-MCTS1 and EIF2D we most strongly upregulated (Suppl. Fig. 6). The occupancy of one of ATF4’s uORFs was reduced in the tumor samples from our dataset, though the change in uORF/CDS ratio between T and NT tissue was not statistically significant (Suppl. Fig. 6). Also highly upregulated in the tumors was EIF5A2, a translation factor broadly implicated in metastasis [98], including in HCC [99]. Its knockdown sensitizes HCC cells to doxorubicin [100]. tRNA synthases as well as components of the elongation factor 1 that delivers tRNAs to the ribosome were also generally upregulated (Suppl. Fig. 6).

We also identified so far unannotated ORFs showing 3-nucleotide periodicity of RPF coverage downstream of annotated coding regions. These could be regions of stop codon readthrough or dORFs. The abundance of readthrough isoforms, estimated from the ribosome densities on CDS and 3’UTRs, is in the range of a few percentages relative to the canonical ORF, consistent with prior estimates [101,102]. The detection of RPF-supported readthrough peptides in proteomics datasets (Suppl. Fig. 5) is further indication that such isoforms are not entirely uncommon. Potential functions of dORF translation are currently debated. Recent studies identified an abundance of downstream cleavage products of alternative polyadenylation (APA) that appear to undergo translation [103], raising the possibility that some of reported dORFs are in fact the CDSs of these APA-derived transcripts. However, conclusive evidence of downstream products of APA undergoing translation is still lacking. Relatively few dORFs identified in our data are supported by mass spectra even when analyzing extensive databases such as PRIDE [65]. This suggests that the dORF-derived peptides are not relevant per se, which is also consistent with their lack of evolutionary conservation [104]. By comparing the translation efficiency of mRNAs containing or not dORFs, it has been previously proposed that dORFs promote the translation of the corresponding CDS [104]. Our analysis of the dORF/CDS ratio and the CDS TE in tumor and non-tumor liver samples did not reveal statistically supported regulatory interactions between dORFs and CDSs (Fig. 4B-C).

Finally, we have identified numerous RNAs that are still annotated as non-coding in the NONCODE database [59], despite having robust evidence of translation, both in our data and in proteomics datasets. In the non-viral background, the TE of lncRNA-derived ORFs is increased, on average (Fig. 5F), counteracting to some extent the transcriptional downregulation (Fig. 5E). In contrast, in the viral background, there is stronger concordance of TO and mRNA level changes (Fig. 5D and a bias for decreased TE (Fig. 5F). Thus, if lncRNAs generate neoantigens to provide targets for immunotherapy [105], the neoantigen load may be different in HCC of viral and steatohepatitis origin leading to dissimilar responses to immunotherapies [106].

Our study thus reveals the landscape of mRNA translation in steatohepatitis-associated HCC and identifies genes that are regulated at transcriptional or translational level. We uncovered numerous translated regions – for which the term translon has been recently coined [107] – including in RNAs currently annotated as non-coding, some yielding peptides that can be reproducibly identified in mass spectrometry datasets. We provide evidence for a CDS-translation-regulatory function of some uORFs in HCC, though uORFs only explain a relatively small fraction of TE changes. Overall, translational control appears to be more prominent in HCC arising on the background of metabolic disease, affecting hundreds of target genes.

## Methods

### Samples collection and histopathological assessment

Samples were collected from patients undergoing surgery at the University Center for Gastrointestinal and Liver Disease (Clarunis), Basel, Switzerland. Written informed consent was obtained from all patients. The study was performed in accordance with the Helsinki Declaration and approved by the ethics committee (Ethics Committee of Basel, EKBB, no. 2019-02118). Histopathologic characterization was performed as per standard of care at the Institute of Pathology Basel from certified pathologists.

### Tissue lysis and sample preparation

Tissues (20-40 mg) were lysed under cryogenic conditions with 100 µl per 10 mg of tissue of lysis buffer (20 mM Tris-HCl pH 7.5, 100 mM NaCl, 10 mM MgCl2, 1% Triton X-100, 0.5% sodium deoxycholate, 1 mM dithiothreitol (DTT), 100 µg/ml cycloheximide, 0.8 U/µl RNasin plus RNase inhibitor (Promega), 0.04 U/µl Turbo DNase (Invitrogen), and EDTA-free protease inhibitor cocktail (Roche)) on an automatic freeze mill (SPEX Sample prep) using the following program: Precool: 2 min, Run: 2 min, Cool: 2 min, Cycles: 2, CPS: 5. Pulverized samples were transferred to a pre-chilled 50 ml falcon tube and incubated with continuous rotation on an overhead shaker (15 rpm) for 60 min at 4°C. Lysates were then passed through a 23G needle for 10 times on ice and kept with continuous rotation on an overhead shaker (15 rpm) at 4°C for 5 min. Lysates were centrifuged at 3’000 xg for 3 min at 4°C, supernatants were collected and centrifuged again at 10’000 xg for 5 min at 4°C. RNA concentration was measured using a Nanodrop 2000. From these clarified lysates, the translatome (ribosome profiling) and the proteome (proteomics) were analyzed.

### Ribosome profiling

RNA concentration from clarified lysates was measured using a Nanodrop 2000 spectrophotometer (ThermoFisher Scientific). Ribosome profiling was performed as described in [87] from equivalents to 4-6 OD_260_ of lysates treated with 8.25 U RNase I (Invitrogen) per unit of OD_260_ for 45 min at 22°C in a thermomixer (1’000 rpm). Libraries were prepared in a ligation-free manner with the SMARTer smRNA-Seq Kit for Illumina (Takara) and sequenced on the Illumina NextSeq 500 sequencer in the Genomics Facility Basel (Department of Biosystems Science and Engineering (D-BSSE), ETH Zürich).

### Analysis of ribosome profiling data

Single-end reads from .fastq files were trimmed with fastx_clipper from FASTX-Toolkit version 0.0.14 with parameters ‘-a (3’adapter) AAAAAAAAAA, -l (minimum-length) 20, -c (discards non-clipped sequences) and -n (discards sequences with unknown (N) nucleotides)’. The trimmed reads were further trimmed with fastq_quality_trimmer from the same toolkit with -t (minimum quality) 20, -Q (quality type) 33. Then the trimmed reads were filtered with fastq_quality_filter from the same toolkit for read quality with the following parameters: ‘-q (minimum quality) 20, -p (minimum percent of bases that must have [-q] quality) 90’, -l (minimum-length) 20. These reads were first aligned to ribosomal RNA (rRNA) sequences obtained from *Homo sapiens* ribosomal DNA (rDNA), complete repeating unit (https://www.ncbi.nlm.nih.gov/nuccore/U13369) using Segemehl 15 version 0.2.0. The reads that did not map to rDNA were then aligned to the longest coding transcripts for each gene identified from Homo sapiens GRCh38–hg38 genome assembly, Ensembl 110 annotation using Segemehl. The uniquely mapped reads from this alignment were used for downstream analysis. Reads were further separated into groups by read lengths. In each such group, all plausible nucleotide distances from 5’end of read to a position of the 1st nucleotide of a presumable ribosomal P-site were compared in terms of compatibility with 1st nucleotide positions of annotated codons across genes. Then, the distance giving highest compatibility was selected as characteristic for a particular read length. P-site positions in reads were further inferred by shifting the position of 5’ends using that characteristic distance. Lastly, reads with determined P-sites from different read length groups were pooled together for every sample. Counts of reads per translation unit per sample are available from Zenodo with DOI: 10.5281/zenodo.18339128.

### Identification of non-canonical open reading frames (ncORFs) and stop-codon readthrough isoforms (SCRTs)

To identify non-canonical translation products (u/dORF and lncRNA-derived peptides and SCRT isoforms, altogether referred to as ncORFs), we employed an *in-house* script (see Code availability section) that first identifies all alternative start and stop codons on all possible frames of untranslated regions (UTRs) of all protein coding transcripts and lncRNAs, based on the input .gtf annotation file and genome sequence. It then quantifies the coverage from inferred P-site locations for all three reading frames within each region. If the number of inferred P-site positions along a candidate ncORF was smaller than 5, the ncORF was rejected. Then we checked the periodicity of RPF coverage by the non-parametric Wilcoxon rank-sum test and accept ncORFs and SCRTs if the *in-frame* coverage is statistically higher than the *alternative* 2 frames (combined p-value by Fisher’s combined probability test < 0.01, for comparisons between *in-frame* and 2 *alternative* frames). We removed any ncORF that overlaps with any annotated transcript isoforms in Ensembl 110 annotation. For ncORFs and SCRT that pass all the filters above, we checked for comprehensive coverage (at least 1 RPF) across all samples of NT or T condition. For uORFs, dORFs and SCRT, we applied this filter for non-viral samples. For lncRNA-derived ncORFs, we applied it separately for viral and non-viral disease backgrounds and then kept ncORFs that satisfy the comprehensive coverage filter in either of the disease backgrounds. For accepted ncORFs and SCRTs we generated the corresponding peptide sequences. Finally, we append all the possible non-canonical peptides to the *homo sapiens* Uniprot [108] protein database for further use in the analysis of LC-MS data.

### Analysis of differential expression, protein output, translational efficiency and differential ncORF/CDS ratio

Differential expression (based on RNA-seq reads) and differential translational output (based on ribosome profiling reads) analyses were performed using the DESeq2 R package [42] version 1.46.0 with default parameters. The deltaTE [43] procedure was applied for differential translation efficiency analysis using only ribosome profiling reads mapped to coding sequence (CDS) regions obtained from the GRCh38–hg38 genome assembly, Ensembl 110 annotation. For differential protein output and differential translation efficiency analysis for lncRNAs, the same procedure above was followed after identifying translated ORFs on lncRNAs and treating them as CDS regions. For differential analysis of u/dORFs, we employed the deltaTE [43] procedure by using condition (c) component in the model as T/NT, sequencing type (s) component as uORF/CDS counts and followed the same procedure as in differential translation efficiency analysis.

### Gene Set Enrichment (GSEA) and Gene Ontology (GO) analysis

The ClusterProfiler [109] R package version 4.14.6 was used for all the GO term analyses reported in this study. ComplexHeatmap [110] version 2.22.0 and circlize [110,111] version 0.4.16 R packages were used to construct the heatmaps. For GSEA, the fgsea R package [112] version 1.32.0) was used, with permutation parameter nperm = 100’000.

### Processing and analysis of the RNA sequencing data from the TCGA project

Genomic alignments (BAM format) of short paired-end reads from 371 primary tumor and 50 non-tumor RNA-seq samples of the TCGA-LIHC project (The Cancer Genome Atlas, liver hepatocellular carcinoma) were obtained from the GDC portal (accession number phs000178.v11.p8). To evaluate the abundance of transcripts supporting the expression of CDSs and ncORFs, their respective transcriptomic coordinates were converted to genomic coordinates with a custom python script. More precisely, each CDS and ncORF was represented as a unique “gene” element with a single transcript and specified exons, represented in the GENCODE GTF annotation format [113]. This GTF file was used as input to the FeatureCounts package [114] to obtain raw fragment counts per gene across samples, with the following parameters: featureCounts -p --countReadPairs -O --fraction -Q 255 -s 0 -B -C -P -d 0 -D 10000000000000 -t exon -g gene_id.

The ISMARA web server [115] requires unmapped reads in .fastq format as input. Thus, we first converted the uniquely mapped reads from BAM files to .fastq using Samtools v. 1.18 [116][117]. Briefly, ISMARA uses a Bayesian procedure to model log_2_-transformed TPM expression levels of all mRNAs in a sample as a linear function of the transcription factor binding sites in the corresponding promoter sequences. The estimated coefficients of the linear model are the *motif activities* in a sample, where the motifs represent the sequence specificity of the transcription factors. An increased motif activity is inferred when the target genes of the respective transcription factor show relative increase in their TPM values. TPM values associated with every promoter are centered around zero across all samples before analysis, therefore across-sample average motif activity is always zero for all motifs. The explanatory power of any given regulatory motif relative to all other motifs across all samples of the data set is represented by its z-score.

### Classification of TCGA data

To split the patient samples into viral and non-viral categories, we used the hepatocarcinoma risk factor information from metadata provided by the TCGA project. Patients with risk factors *Hepatitis B, Hepatitis C, Hepatitis C|Other, Hepatitis B|Hepatitis C, Hepatitis B|Other* are labeled as viral samples (12 NT, 110 T). Patients with risk factors *Alcohol consumption|Nonalcoholic Fatty Liver Disease, Alcohol consumption, Nonalcoholic Fatty Liver Disease, Alcohol consumption|Other, Alcohol consumption|Nonalcoholic Fatty Liver Disease|Other, Nonalcoholic Fatty Liver Disease|Other* are labelled as non-viral samples (11 NT, 90 T).

### Mass spectrometry-based proteome analysis

Protein samples (see “Tissue lysis and sample preparation” section) were incubated for 10 min at 95°C. After cooling, a 1 M iodoacetamide solution was added to reach a final concentration of 20 mM and the cysteine residues alkylated for 30 min at 25°C in the dark. Subsequently, the protein concentration was determined by fluorometric analysis of tryptophane content as described [118]. Sample aliquots containing 50 μg of total proteins were then purified and digested with the S-Trap cartridges (Protifi, Fairport NY, USA) following the manufacturer’s instructions. Resulting peptides were dried under vacuum and stored at −20°C until LC-MS analysis.

Peptides were re-suspended in 0.1% aqueous formic acid and 0.2 µg of peptides subjected to LC–MS/MS analysis using an Orbitrap Exploris 480 MassSpectrometer fitted with an Vanquish Neo (both Thermo Fisher Scientific) and a custom-made column heater set to 60°C. Peptides were resolved using a RP-HPLC column (75 μm × 30 cm) packed in-house with C18 resin (ReproSil-Pur C18–AQ, 1.9 μm resin; Dr. Maisch GmbH) at a flow rate of 0.2 μl/min. Separation of peptides was achieved using the following gradient: Proteome analysis: 4% buffer B to 10% in 5 min, to 35% buffer B in 45 min, to 50% buffer in 10 min. Buffer A was 0.1% formic acid in water and buffer B was 80% acetonitrile, 0.1% formic acid in water. The mass spectrometer was operated in DIA acquisition mode with a total cycle time not exceeding approximately 3 s. For MS1, the following parameters were set: Resolution: 120,000 FWHM (at 200 m/z); Scan Range: 350-1400 m/z; Injection time: 25 ms; Normalized AGC Target: 300%. MS2 (SWATH) scans were acquired using the following parameters: Isolation Window: 10 m/z; HCD Collision Energy (normalized): 28%; Normalized AGC target: 1000%; Resolution: 15,000 FWHM (at 200 m/z); Precursor Mass Range: 400 – 900 m/z; Max. Fill Time: 22 ms, DataType: Centroid. In total 49 DIA (MS2) mass windows per MS cycle followed by the one MS1 scan.

### Analysis of LC-MS-based proteomics data

The acquired raw files were first converted to indexed mzML format using ThermoRawFileParser [119] and then searched against a protein database containing sequences of the SwissProt Release 2024 [108,119] entries of *Homo sapiens* (in total 20’420 protein sequences) along with commonly observed contaminants using using the workflow for data-independent acquisition (downloaded from https://github.com/Nesvilab/FragPipe/releases) of FragPipe [120] version 21.0.0. For non-canonical peptide identification, the peptide-spectrum, ion and peptide matches were filtered to 1% FDR level. IBAQ values are calculated as the intensity of a protein estimated by DIA-NN approach [121] divided by the number of theoretically observable tryptic peptides for that protein.

All mass spectrometry proteomics data associated with this manuscript have been deposited to the ProteomicsXchange consortium via MassIVE (https://massive.ucsd.edu) with the accession number MSV000099492 (Dataset is currently private. Reviewers can access the dataset following the instructions on the website. Username: MSV000099492_reviewer, Password: PCF). Details on the correspondence between sample ID (MassIVE) and biopsy name are in Suppl. Table 1.

## Abbreviations

HCC: Hepatocellular carcinoma
ORF: open reading frame
CDS: coding region
ncORF: non-canonical ORF
lncRNA: long non-coding RNA
HBV: hepatitis B virus
HCV: hepatitis C virus
TCGA: the cancer genome atlas
SH: steatohepatitis
ASH: alcoholic SH
NASH: non-alcoholic SH
NT: non-tumor
T: tumor
RPF: ribosome-protected fragment
TO: translational output
TE: translation efficiency
TPM: transcripts per million
PCA: principal component analysis
TF: transcription factor
ISMARA: integrated motif activity response analysis
UTR: untranslated region
RT: readthrough

## Acknowledgements

This work was supported in part by a grant from the Helmut Horten Foundation and by the Swiss National Science Foundation grant # 310030_204517 to M.Z. The results shown here are in part based upon data generated by the TCGA Research Network: https://www.cancer.gov/tcga.

## Contributions

M.Z. conceived the study. A.G. and M.P. performed ribosome profiling and proteomics experiments. M.A., A.M., N.S., and M.Z. analyzed all data. A.S. carried out mass spectrometry experiments. M.C-L. and S.P. provided biopsies. N.M. provided advice on ribosome profiling experiments. M.Z. wrote the manuscript with help from A.G., M.A., A.M., N.S., M.C-L, and S.P. Funding was obtained by M.Z. All authors have read and approved the manuscript.

## Code availability

The custom scripts for analyzing the ribosome profiling, RNA-sequencing and proteomics data are deposited on Zenodo with DOI: 10.5281/zenodo.18339128.

## Supplementary Figures

**Figure S1.**
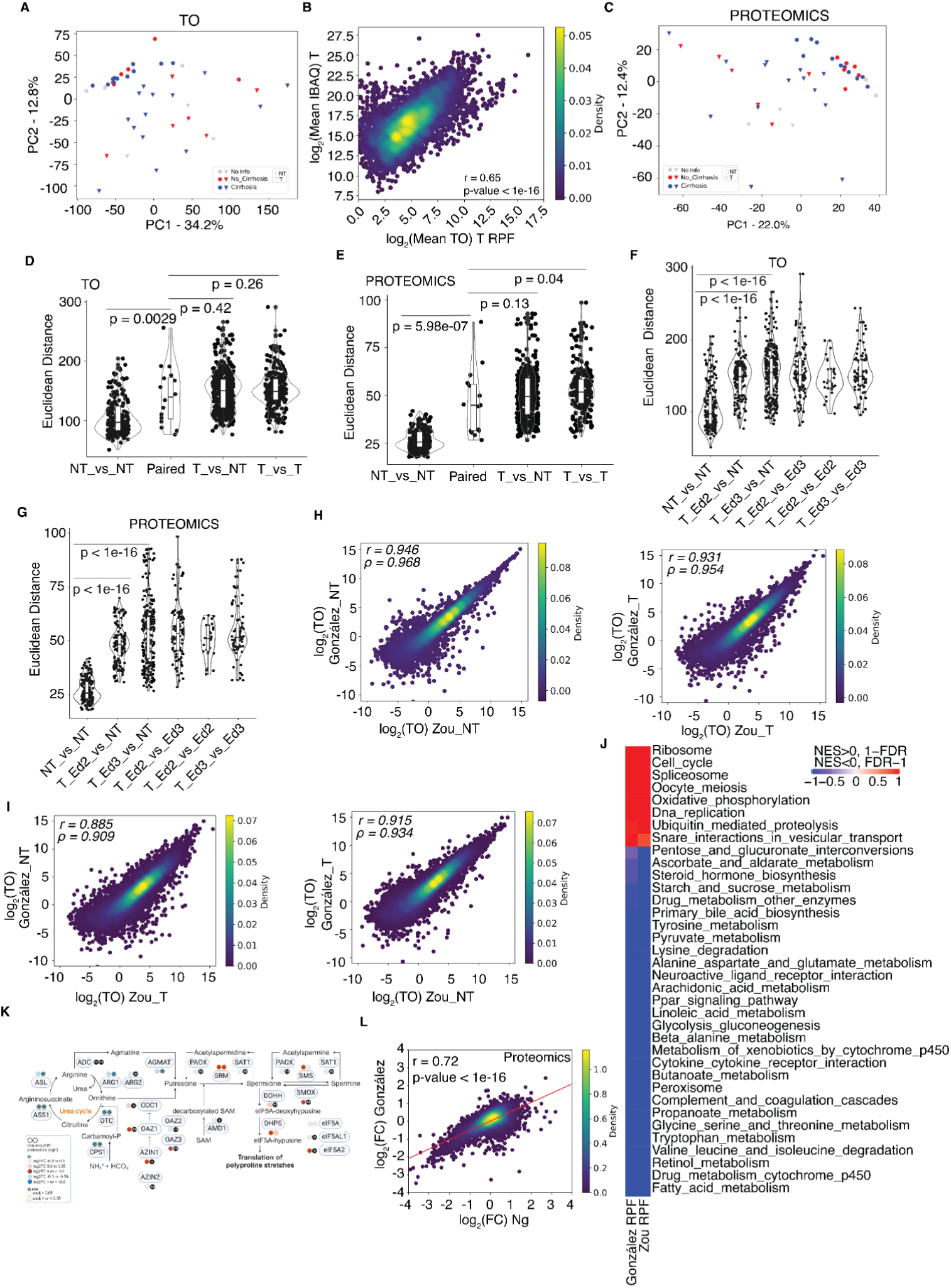
Comparative PCA, correlation, and pathway analyses across ribosome profiling and proteomics data. **A.** Principal component analysis (PCA) of TO data labeled by presence/absence of cirrhosis. **B.** Correlation of TO (RPF-based TPM) with the protein intensity calculated as Intensity-Based Absolute Quantification (iBAQ) signal for tumor samples. **C.** As in A, based on proteomics data. **D.** Euclidean distances between TOs of individual genes in pairs of all NT, T-NT, T-T and paired T-NT samples from viral/non-viral disease background groups. **E.** Same as D, for protein intensity values. **F.** Euclidean distances between TO values of individual genes in pairs of NT and T samples grouped by Edmondson’s grade. **G.** Same as F, from protein intensity values of individual genes in pairs of NT and T samples grouped by Edmondson’s grade. **H.** Correlation of average RPF TPM values from NT (panel on left) samples from this study and Zou study [30] and same for T samples (panel on right). **I.** Same as H, for NT samples from this study and T from Zou study (panel on left) and same for NT samples from Zou study, T samples from this study (panel on right). **J.** GSEA analysis using KEGG pathways on RPF data in this study and Zou study, shown pathways have adjusted p-value < 0.001. **K.** Urea cycle, arginine biosynthesis and polyamine synthesis pathways colored by log_2_FC from ribosome profiling (circles on the left) and proteomics data (circles on the right). Created with BioRender, design based on [36]. **L.** Correlation of log_2_FC values (T. vs. NT) using proteomics data, Ng study [8] on x-axis, this study on y-axis.

**Figure S2.**
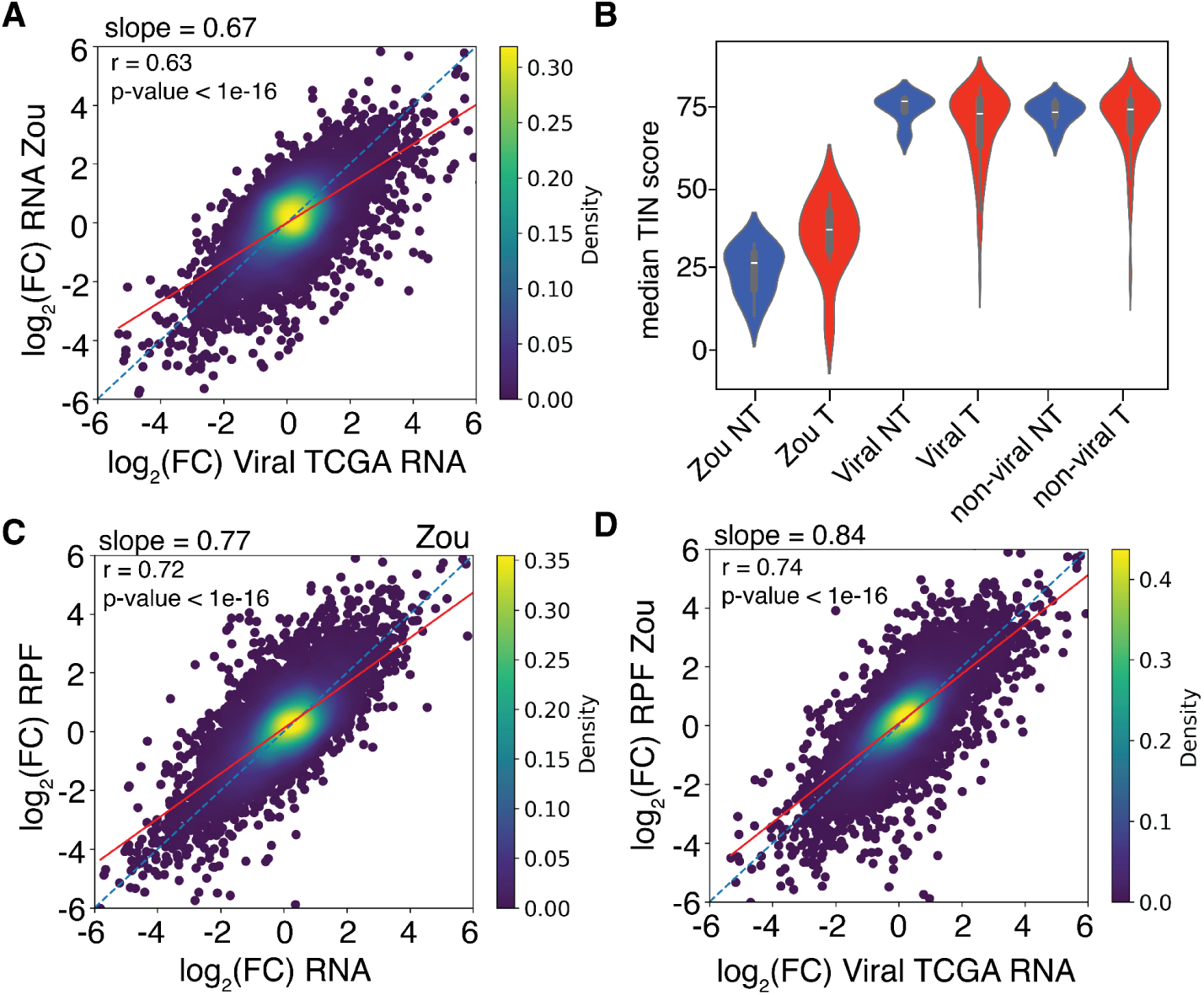
Cross-dataset correlation of transcriptome changes. **A.** Correlation of mRNA level changes in T vs. NT in TCGA viral samples (x-axis), and mRNA level changes in the Zou dataset. **B.** Median TIN scores across the transcriptome for NT and T samples from Zou dataset and for NT and T samples from TCGA samples with viral and non-viral backgrounds. **C.** Correlation of mRNA level changes in T vs. NT samples in mRNA level (x-axis) and RPF level (y-axis) changes in the Zou dataset. The diagonal line (orange, dashed) corresponds to changes in translational output that are entirely explained by changes in mRNA levels. **D.** Same as A, for mRNA level changes in T vs. NT in TCGA viral samples (x-axis), and mRNA level changes in T vs. NT in the Zou dataset.

**Figure S3.**
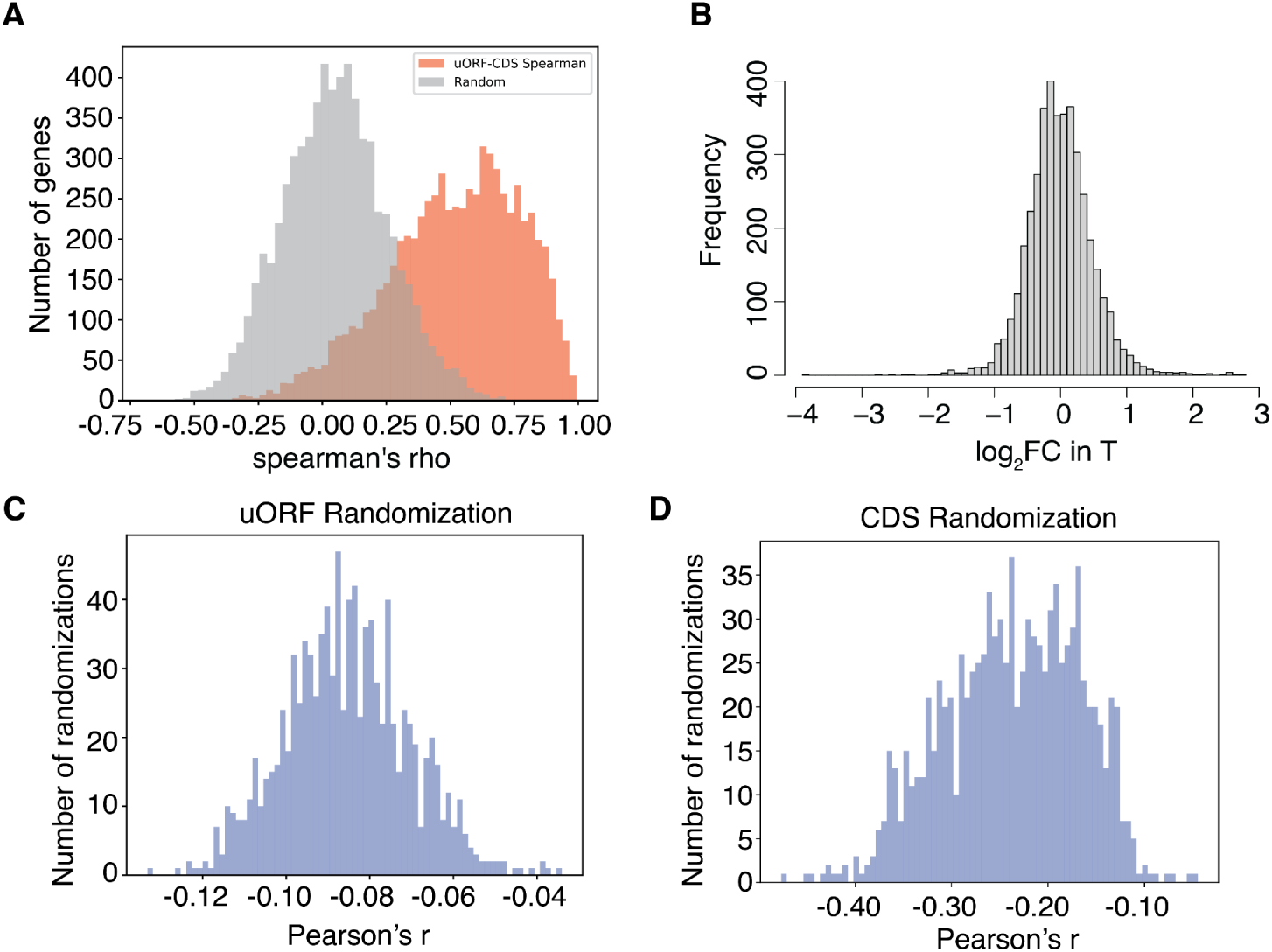
Analysis of uORF-CDS coordination. **A.** Distribution of Spearman correlation coefficients of uORF and corresponding CDS TO values for individual genes across all 43 samples (orange bars). In gray are the Spearman correlation coefficients when the uORF TPMs were permuted (10000 randomizations). **B.** Distribution of log_2_FC in the uORF/CDS ratio in non-viral-association T relative to NT samples. **C.** Distribution of Pearson’s correlation coefficient *r* of the uORF/CDS ratio changes with the TE changes of the corresponding CDS, calculated after shuffling the uORF RPF counts (1000 randomizations). The analysis was done for samples with no virus association. **D.** Same as C, when CDS RPF were shuffled before calculating the uORF/CDS ratios and the TEs.

**Figure S4.**
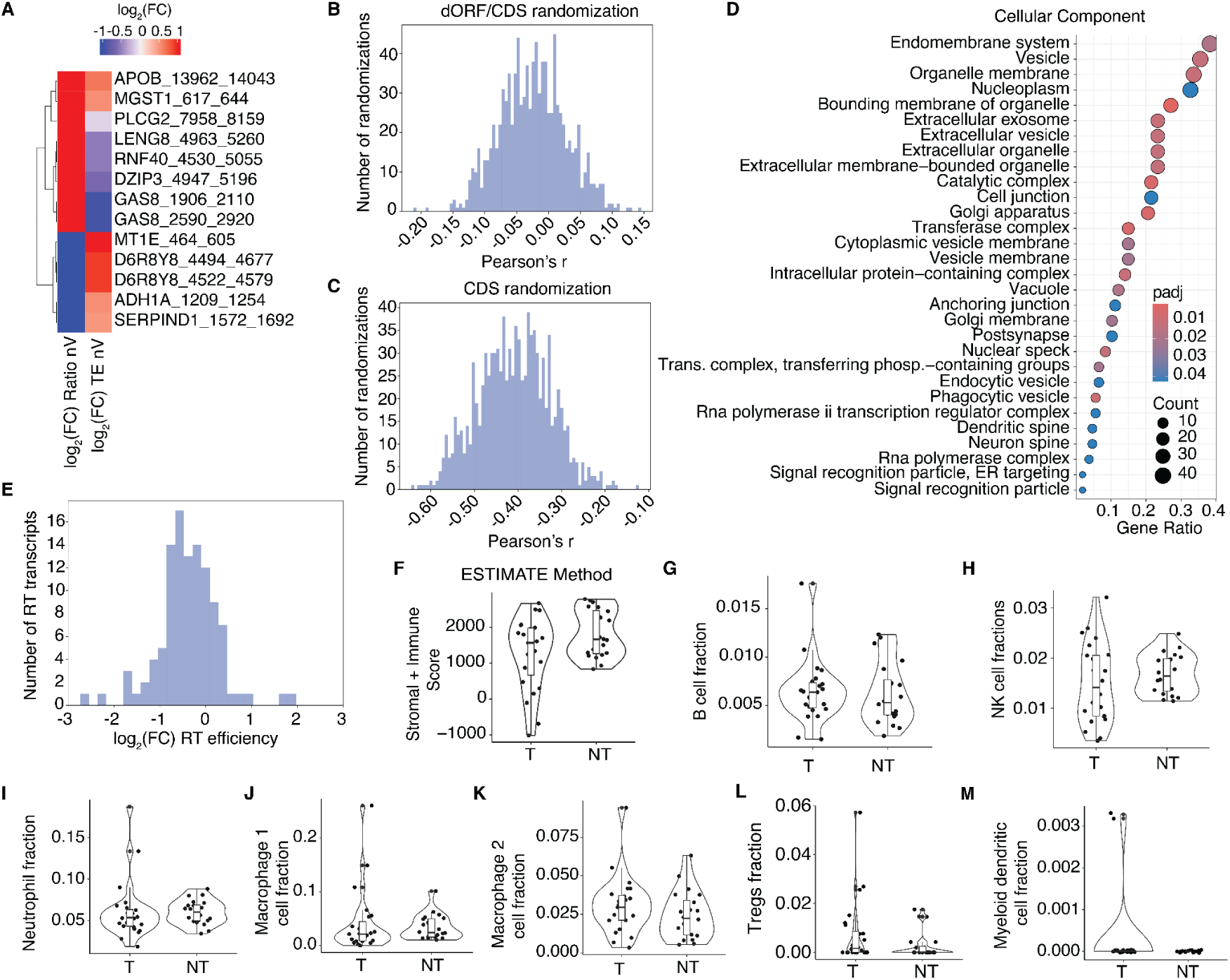
Analysis of dORF/CDS ratios, readthrough efficiency, and immune signatures. **A.** Genes with significant dORF/CDS ratio (p-value<0.01, according to customized DESeq2 analysis) and the corresponding TE changes in T vs NT in non-viral backgrounds. **B.** Distribution of Pearson’s *r* values for the correlation of dORF/CDS ratio and the corresponding TE of CDS, where the dORF RPF counts were shuffled before calculating the dORF/CDS ratios (1000 randomizations) for non-viral samples. **C.** Same as C, when CDS RPF were shuffled before calculating the uORF/CDS ratios and the TEs. **D.** Cellular Component (CC) GO terms overrepresentation analysis for genes with readthrough (RT) evidence. **E.** log2FC of RT efficiency in T vs NT samples in non-viral backgrounds. **F.** Stromal and immune score for T and NT samples estimated by ESTIMATE method [122]. **G.** B-cell fraction in T and NT samples estimated by quanTIseq method [123]. **H.** Same as B, for NK cells, for neutrophil **(I)**, for macrophage 1 **(J),** for macrophage 2 **(K),** for tregs **(L)** and for myeloid dendritic cells **(M).**

**Figure S5.**
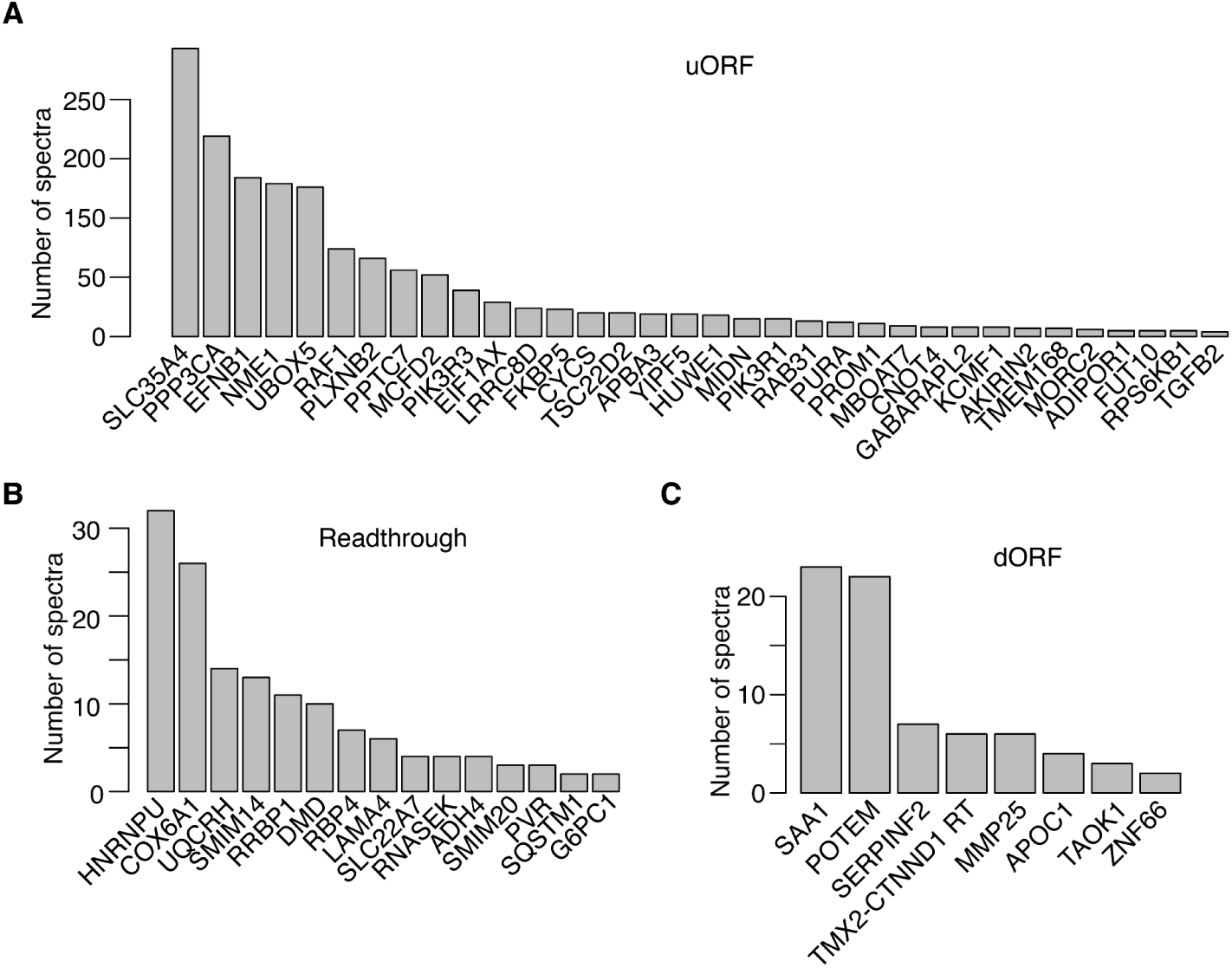
Proteomics evidence for ncORFs. **A.** uORF derived peptides, with spectral evidence from at least 3 datasets. **B.** readthrough-isoforms from at least 2 datasets. **C.** dORF derived peptides from at least 2 datasets.

**Figure S6.**
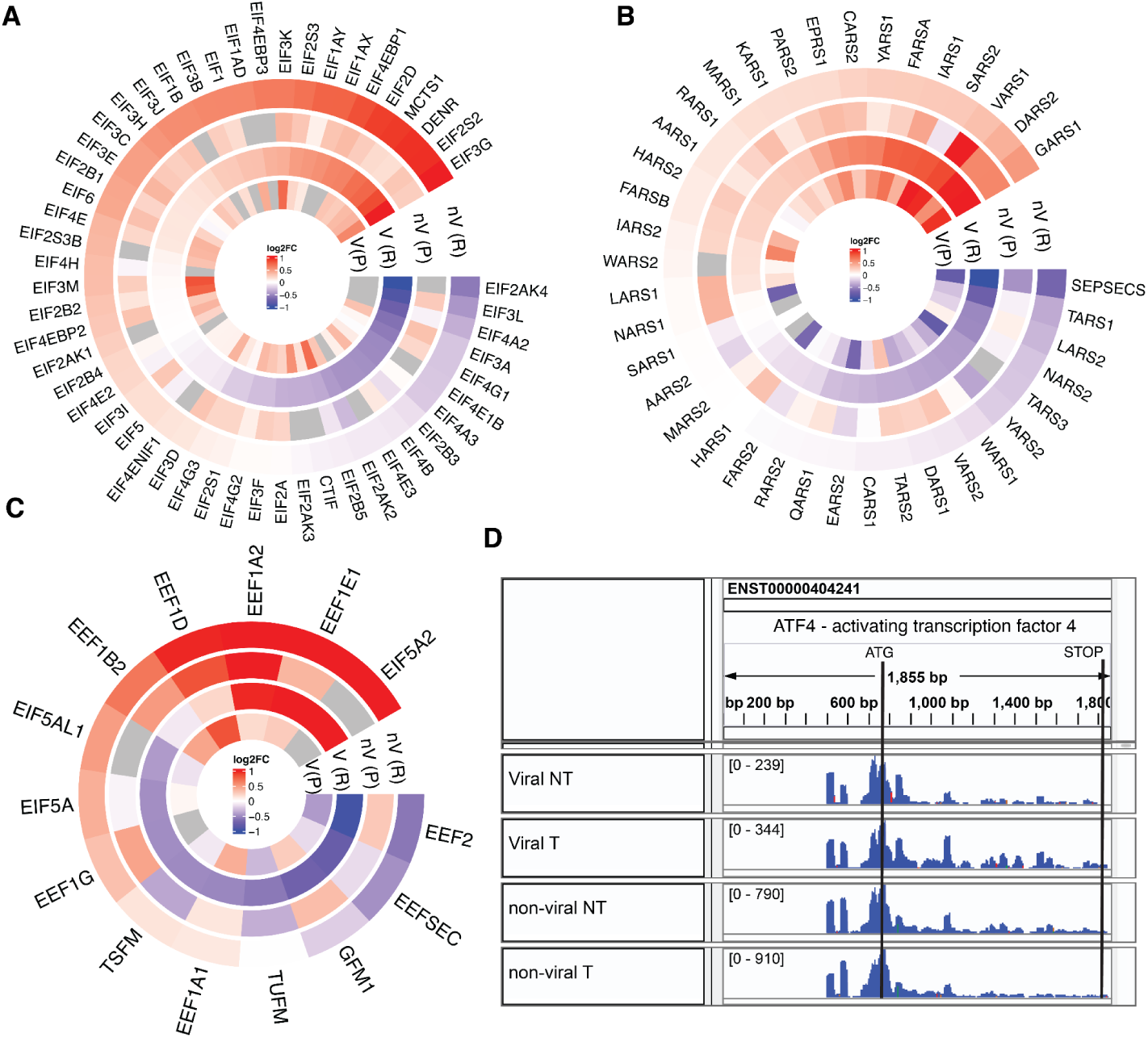
Translation factors are generally upregulated at RPF level in tumor samples relative to non-tumor. **A.** Translation initiation factors. **B.** tRNA synthases. **C.** Translation elongation factors. Shown are heatmaps of log_2_ fold changes in tumor relative to non-tumor, the two heatmaps showing changes at RPF level (R) and protein level (P) for viral and non-viral backgrounds. **D.** IGV snapshot of ATF4 transcript RPF coverage.

